# High-throughput identification of FLT3 wild-type and mutant kinase substrate preferences and application to design of sensitive *in vitro* kinase assay substrates

**DOI:** 10.1101/457689

**Authors:** Minervo Perez, John Blankenhorn, Kevin J. Murray, Laurie L. Parker

**Affiliations:** University of Minnesota, Department of Biochemistry, Molecular Biology and Biophysics, 420 Washington Avenue SE, Minneapolis, MN 55455; Purdue University, Department of Medicinal Chemistry and Molecular Pharmacology, 201 S. University Street, West Lafayette, IN 47907; University of Minnesota, Department of Veterinary Population Medicine, 319 15th Avenue South East, Minneapolis, MN 55455

**Keywords:** Acute Myeloid Leukemia, FLT3, Mass spectrometry, Phosphoproteomics, Kinase Substrate Prediction, KINATEST-ID Pipeline, Kinase Assay-Linked Phosphoproteomics, Assay Development

## Abstract

Acute myeloid leukemia (AML) is an aggressive disease that is characterized by abnormal increase of immature myeloblasts in blood and bone marrow. The FLT3 receptor tyrosine kinase plays an integral role in haematopoiesis, and one third of AML diagnoses exhibit gain-of-function mutations in FLT3, with the juxtamembrane domain internal tandem duplication (ITD) and the kinase domain D835Y variants observed most frequently. Few FLT3 substrates or phosphorylation sites are known, which limits insight into FLT3’s substrate preferences and makes assay design particularly challenging. We applied *in vitro* phosphorylation of a cell lysate digest (adaptation of the Kinase Assay Linked with Phosphoproteomics (KALIP) technique and similar methods) for high-throughput identification of substrates for three FLT3 variants (wild-type, ITD mutant, and D835Y mutant). Incorporation of identified substrate sequences as input into the KINATEST-ID substrate preference analysis and assay development pipeline facilitated the design of several peptide substrates that are phosphorylated efficiently by all three FLT3 kinase variants. These substrates could be used in assays to identify new FLT3 inhibitors that overcome resistant mutations to improve FLT3-positive AML treatment.

## Introduction

Acute myeloid leukemia (AML) is an aggressive cancer with a diverse genetic landscape. The FLT3 gene encodes for a receptor tyrosine kinase (FLT3) that regulates hematopoiesis and perturbations to its signaling pathways appear to promote AML disease progression. In fact, FLT3 is implicated as a major factor in AML relapse.^1^ Thirty percent of AML cases have mutations to FLT3 that lead the kinase to be constitutively active,^2,3^ most commonly to the juxtamembrane domain and the kinase domain.^2,4,5^ Internal tandem duplication (FLT3-ITD) in the juxtamembrane or the first tyrosine kinase domain (TKD) occurs when a segment is duplicated (head to tail) leading to the loss of repressive regions in the protein.^6^ A second common mutation is a substitution of aspartic acid 835 to a tyrosine residue (D835Y) in the TKD. Both ITD and TKD mutants can activate and dimerize with the wild type FLT3.^7^ The effects of these mutations on FLT3 signaling are still unclear, but one possibility is that mutant FLT3-TKD and FLT3-ITD activate alternative signaling pathways, or activate standard FLT3 pathways aberrantly, compared to the WT. Mutations to FLT3 are correlated with poor long-term prognosis^8,9^ and while patients with FLT3 mutations achieve similar initial disease remission to those with wild-type FLT3, they have an increased risk for relapse.^2,8,10^ *In vitro* studies show that FLT3-ITD mutant-expressing cell lines are resistant to cytosine arabinoside (the primary AML therapeutic).^8^ These findings prompted the use of a combinatorial approach to AML therapies to include FLT3 tyrosine kinase inhibitors (TKIs), which are frequently initially successful but often lead to FLT3 inhibitor resistance and subsequent disease relapse.

The current FDA approved TKIs used to inhibit FLT3 were not developed specifically to target FLT3.^11–13^ Sorafenib is a type II pan-TKI which is FDA approved for use in combinatorial approaches with AML chemotherapy, but elicits no response in FLT3 variants with tyrosine kinase domain mutations.^8,14–17^ Efforts to develop FLT3 mutant-specific TKIs lead to the discovery of the type II TKI quizartinib, which can inhibit the FLT3-ITD mutant and is currently undergoing phase III clinical trials for AML.^18^ However, quizartinib has no activity against FLT3-TKD point mutations and thus these mutations are the primary mode of quizartinib monotherapy resistance.^18–21^ Quizartinib also has potent activity towards Platelet Derived Growth Factor receptor (PDGFR) and c-KIT kinases, and produces side effects that may be related to their inhibition in patients undergoing a FLT3 TKI regimen.^22,23^ Crenolanib, a TKI designed to target the α and β isoforms of PDGFR, has demonstrated activity against a broad range of FLT3 mutations.^1,24^ Unlike quizartinib, crenolanib does not inhibit c-KIT (the main kinase implicated in undesirable side effects of quizartinib) at safe plasma concentrations, and is undergoing phase II clinical trials in relapsed AML patients with a driver FLT3 mutation (NCT01657682).^22,25^ However, recent reports have shown that secondary point mutations within the kinase domain of FLT3 can reduce crenolanib’s clinical efficacy that suggest it is only a matter of time until crenolanib resistant mutations are found in a clinical setting.^22,25^

The complex abnormality landscape of AML reduces the possibility that a single FLT3 TKI would be a viable monotherapy for AML. Although crenolanib is a promising TKI, efficient development of new inhibitors will require better assays than those currently available, and adaptable strategies that effectively screen inhibitors to target mutant forms of FLT3 are especially needed.^18^ Since very little is known about FLT3 substrate preferences, there are few options available when designing FLT3 activity assays. The current activity tests are limited by inefficient phosphorylation activity, and/or their phosphorylation by the mutant variants has not been characterized. In this manuscript, we describe the development of several novel and efficient peptide substrates for FLT3 and two clinically-significant mutant variants (the ITD and D835Y mutants). We adapted the “Kinase Assay Linked with Phosphoproteomics” (KALIP)^26,27^ strategy (from the Tao lab) to perform high-throughput determination of FLT3’s preferred peptide substrate motif in a manner similar to other previously reported methods (e.g. Kettenbach *et al* from the Gerber group).^28^ In these approaches, a cell lysate digest is stripped of endogenous phosphorylation and used in a kinase reaction as a pseudo-”library” of peptides to determine kinase substrate preferences by identifying phosphorylated sequences via enrichment and mass spectrometry (ideal for high-throughput analysis of many substrates simultaneously without requiring radioactivity or other labeling).^29,30^ We then used the identified substrate preferences to rationally design a panel of candidate peptides incorporating key sequence features predicted to make them favorable for phosphorylation by the FLT3 kinase variants, following our previously reported substrate development pipeline KINATEST-ID.^31^ We demonstrated that these substrates enable efficient inhibitor screening for all three forms of FLT3. These peptides could be used in many different types of drug discovery settings to more rapidly and efficiently screen for and validate FLT3 inhibition.

## Materials and Methods

### Cell Culture and Endogenous Peptide Sample Preparation

KG-1 cells (ATCC) were maintained in IMDM media (Gibco) supplemented with 10% heat inactivated fetal bovine serum (FBS), 1% penicillin/streptomycin in 5% CO_2_ at 37 °C. KG-1 cells were washed with 30 mLs of phosphate buffered saline (PBS) 5 times. The cells were then pelleted at 1,500 RPM for 5 minutes and lysed with buffer containing 8 M urea, 0.1 M ammonium bicarbonate pH 8.5, 20% acetonitrile (ACN), 20 mM dithiothreitol (DTT), and 1X Pierce Phosphatase Inhibitor tablet (Roche) pH 8.0. Lysed cells were incubated on ice for 15 minutes and then were subjected to probe sonication to shear the DNA. Lysates were treated with 40 mM iodoacetamide and incubated at room temperature (protected from light) for 60 minutes. Samples were then centrifuged at 15,000 RPM for 30 minutes to remove cellular debris. Urea concentration was diluted to 1.5 M using 50 mM ammonium bicarbonate buffer (pH 8.0) and the samples were set up for trypsin digestion at a 1:50 trypsin (ThermoScientific) ratio and incubated at 37 °C overnight. Trypsin digestion was quenched by adding 10% trifluoroacetic acid (TFA) in water to lower the pH below 3. Subsequently, the tryptic digest was desalted using hydrophilic-lipophilic balanced copolymer (HLB) reverse phase cartridges (Waters) and vacuum dried.

### Alkaline Phosphatase Treatment

Samples were reconstituted in alkaline phosphatase dephosphorylation buffer containing 50 mM tris(hydroxymethyl)aminomethane hydrochloride (Tris-HCL), 0.1 mM Ethylenediaminetetraacetic acid (EDTA) at pH 8.5. Alkaline phosphatase (6 U, Roche) were added to each sample followed by incubation for 90 minutes at 37 °C. The reaction was quenched by incubating the samples in 75 °C for 15 minutes (Figure 1).

**Figure 1.**
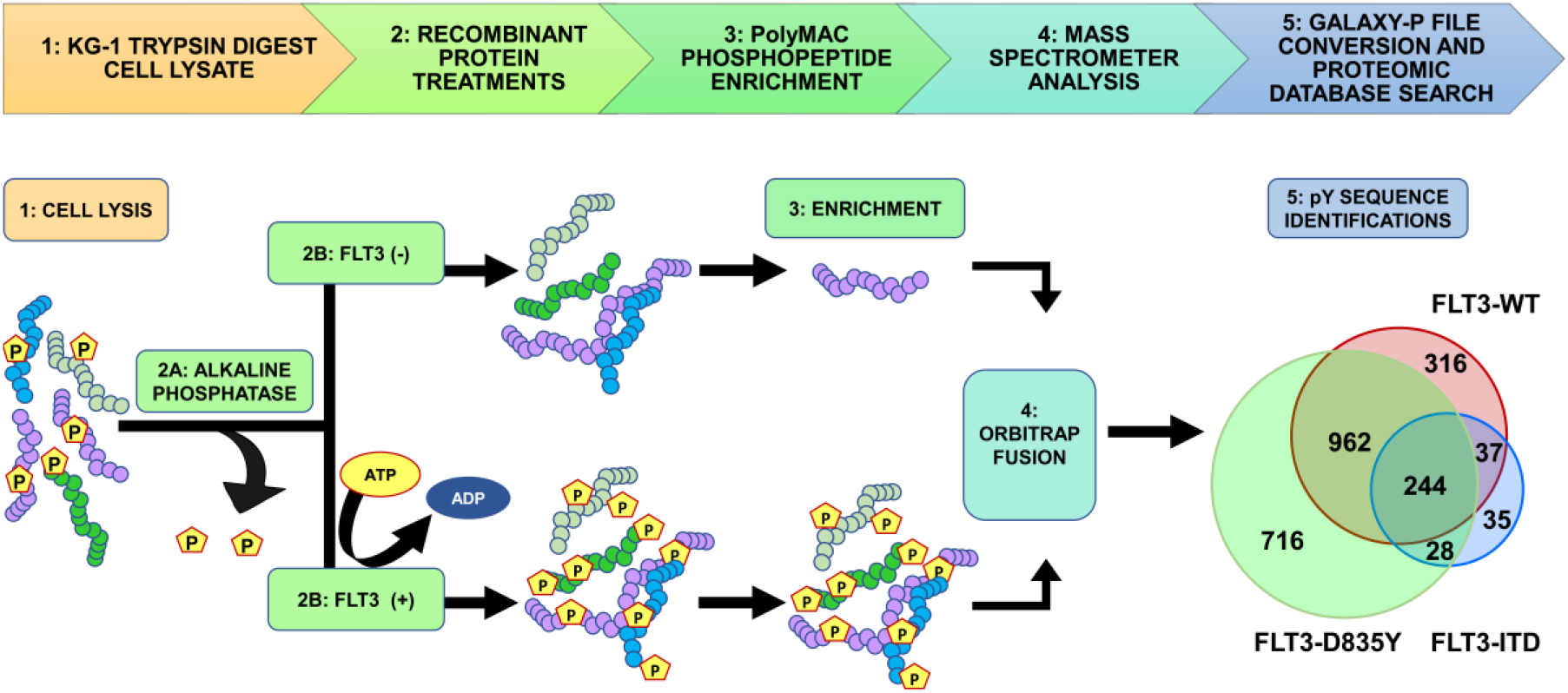
*In vitro* phosphorylation, enrichment and identification of substrate peptides from cell lysate. To identify a larger number of FLT3 kinase substrates than was previously available, we subjected KG-1 cell lysate to trypsin digestion (Step 1). The tryptic digest was then treated with alkaline phosphatase (Step 2A). Subsequently, the sample was split into two equal parts (Step 2B) prior to *in vitro* recombinant FLT3 treatment (FLT3-WT, FLT3-D835Y and FLT3-ITD). The samples were enriched with PolyMAC magnetic beads (Step 3) and an aliquot was analyzed on an Orbitrap-Fusion mass spectrometer (Step 4). The mass spectrometer files were uploaded to the Galaxy-P proteomics pipeline for file conversion and ProteinPilot database search (Step 5).

### KALIP Recombinant FLT3 Kinase Assays

Recombinant kinases were purchased from EMD Millipore (WT, PN: PV3182; FLT3-D835Y, PN: PV3967; FLT3-ITD, PN: PV6190). The samples were briefly vortexed and aliquoted into two equal parts. Peptide samples were reconstituted in kinase reaction buffer containing 50 mM Tris HCL, pH 7.5, 10 mM MgCl_2_, 1 mM DTT, 1 mM Na_3_VO_4_ and 2 mM adenosine 5’-triphosphate (ATP). The kinase (or water for control) was added last to each sample and incubated for 16 hours (16 H) at 37 °C. The FLT3-WT treatment contained an additional two-hour (2H) time point. The reaction was quenched by bringing up the concentration of TFA to 0.5% and desalted using Oasis HLB 1cc cartridge columns with 30-micron particle size and 30 milligram sorbent.

### PolyMAC Enrichment

The phosphopeptide enrichments were carried out according to manufacturer’s instructions (Tymora Analytical, West Lafayette, IN).^26,27^ The enrichment kit is made up of four components: 1) loading buffer, 2) PolyMAC magnetic beads, 3) wash buffer 1 and 2, and 4) elution buffer. In brief, the dried peptides were re-suspended in Loading Buffer and 100 μL of PolyMAC capture beads were added to the mixture. The phosphopeptide-PolyMAC mixture was mixed at 700 RPM for 30 minutes. Subsequently, the mixture was centrifuged briefly and placed on a magnetic rack to remove the un-phosphorylated peptide solution. The beads were washed twice with wash buffer 1 and rocked for 5 minutes at 700 RPM. The phosphopeptide-PolyMAC complex was placed on the magnetic stand until beads were immobilized by the magnet, and the supernatant was discarded; the process was repeated using wash buffer 2. The phosphopeptides were eluted from the capture beads using 300 μL of elution buffer and then vacuum dried.

### LC-MS/MS Data Acquisition

Samples were reconstituted in 25 μL of mass spectrometry loading buffer (98/2/0.5%; H_2_O/ACN/formic acid (FA)) and centrifuged for 30 minutes at 15,000 RPM. A 20 μL aliquot was transferred to a low binding safe-lock microtube (Eppendorf). A 2.5 μL aliquot was loaded on a ThermoScientific Easy NanoLC LC 1000 system. The reverse phased HPLC peptide separation was performed using a 100 μm inner diameter Picotip emitter column packed in-house with 1.9 μm C18 ReproSil-Pur sorbent. The mobile phase consisted of 0.1% formic acid in ultra-pure water (Solvent A) and 0.1% formic acid in acetonitrile (Solvent B). Samples were run over a linear gradient (2-30% solvent B; 60 minutes) with a flow rate of 200 nL/min into a high resolution Orbitrap Fusion Tribrid Mass Spectrometer, operated using data dependent mode at a resolution of 60,000 with a scan range of 300-1500 m/z. After each round of precursor detection, an MS/MS experiment was triggered on the top 12 most abundant ions using High Collision Dissociation (HCD). The mass analyzer parameters were set between two and seven charge states with a dynamic exclusion time of 15 seconds.

### Data Analysis—Phosphopeptide identification

The Orbitrap Fusion mass spectra files were searched against a merged version of the reviewed human Uniprot database downloaded from uniprot.org (2/27/2017; 20,202 entries) and the cRAP database (common lab contaminants; downloaded from (thegpm.org/crap/) on 2/27/2017) using the Paragon algorithm in the ProteinPilot 5.0 proteomic search engine within the Galaxy-P pipeline to create a Distinct Peptide Report (the output report from ProteinPilot 5.0).^32–34^ Peptide precursor mass tolerance was set at 0.02 Da and MS/MS tolerance was set to 0.1 Da. Proteomic database search parameters included trypsin digestion, urea denaturation, phosphorylation emphasis, iodoacetamide fixed modification to cysteine residues, and variable biological modifications.False discovery rate (FDR) analysis was activated for each individual search. ProteinPilot 5.0 used a reverse database as the decoy to calculate the false discovery rate (FDR) for each independent search^35^ and we set the global 1% FDR score as our cutoff threshold.

#### Data processing and KINATEST-ID substrate candidate prediction

##### Streamlined data processing of LC-MS data as input for KINATEST-ID algorithm substrate design

A series of novel scripts were developed to prepare and analyze the results from KALIP to design potential substrates in the KINATEST-ID platform. These steps and scripts are described in more detail in the supplemental methods, and detailed instructions on running each script in sequence are provided in the supporting information file “kinatestsop.docx”. To extract and reformat the phosphopeptide sequences from the ProteinPilot distinct peptide report, we created the KinaMine program and GUI that extracts all sequences from a ProteinPilot 5.0 (SCIEX) Distinct Peptides Report output file that have phosphorylated tyrosine residues identified at a 99% confidence (1% FDR), and creates “Substrate” and “Substrate Background Frequency (SBF)” files, which contain the observed substrate sequences and the UniProt (uniprot.org) accession numbers and calculated representation of all amino acids for the proteins from which substrate sequences were identified, respectively. We created the “commonality and difference finder.r” script to identify the phosphopeptides from the “substrates” and SBF files that are shared by all of the FLT3 kinase variants, and generated the “SHARED-16H” substrate and SBF files. We extracted the UniProt accession numbers from the SBF lists and used them to download a customized FASTA file from the UniProt website that contained entries only for those protein sequences, and converted that into .csv format using the “FASTAtoCSV” script. We created the “Negative Motif Finder.r” script to extract (and *in silico* trypsin digest) all tyrosine centered sequences present in any of the “background” proteins, and compared them to the substrate list (KinaMINE output) to return the sequences that were not observed in the phosphoproteomics data as a best estimate of “non-substrates.”

##### KINATEST-ID streamlined processing in R

Using Substrates, Substrate Background Frequency, Non-substrate Motifs, and Screener.csv, the scripts “Kinatestpart1.R” and “Kinatestpart2.R” were written to replicate the functionality of the KINATEST-ID workbooks previously described,^31^ including determining over‐ and under-representation of particular amino acids (the *standard deviation table/positional scoring matrix*) and/or side chain chemical properties (the *site selectivity matrix*) at particular positions relative to the tyrosine to define a preferred substrate motif, permutation of that motif into a list of all possible combinations of the preferred amino acids at their given positions (*Generator*), and scoring of those sequences against the targeted kinase’s positional scoring matrix model as well as those for a panel of other off-target kinases (*Screener*). These scripts ultimately create three .csv output files named by the user and containing the following information, respectively (all consistent with steps and components of the original published KINATEST-ID implementation)^31^: 1) the Standard Deviation and the AA Percent Tables; 2) the Site Selectivity Matrix, the Endogenous Probability Matrix (EPM) (which gives the scoring function used to calculate scores for a given sequence via the positional scoring matrix model defined by a given input dataset), the Normalization Score and the MCC Characterization Table; and 3) the list of predicted substrates ranked according to lowest “off-target” kinase scores using the Screener comparisons. For more information about these functions see the previously published description of KINATEST-ID.^31^

### Peptide Synthesis and Purification

Peptides were synthesized using a Protein Technologies SymphonyX synthesizer using 4-methylbenzydrylamine hydrochloride resin (Iris Biotech GMBH). Standard Fmoc-protected amino acid (AA) coupling occurred in the presence of 95 mM HCTU (Iris Biotech GMBH) and 200 mM N-methylmorpholine (Gyros Protein Technologies; S-1L-NMM) over two 20-minute coupling cycles. Fmoc deprotection occurred in the presence 20% piperidine in dimethylformamide (DMF, Iris Biotech GMBH;) over two 5-minute cycles. The peptides were purified to >95% purity by preparative C18 reverse phase HPLC (Agilent 1200 series) over a 5-25% acetonitrile/0.1% TFA and water/0.1% TFA gradient and characterized using HPLC-MS (Agilent 6300 MSD). Peptide substrates were dissolved in a PBS solution containing 5% dimethyl sulfoxide (DMSO). Absorbance measurements at 280 nm wavelength were used to determine the peptide concentration using the Beer-Lambert law (for which peptide extinction coefficients were calculated using Innovagen’s peptide property calculator (https://pepcalc.com/).

### In Vitro Kinase Assays

Recombinant kinases were purchased from SignalChem (FLT3, FLT3-D835Y, FLT3-ITD, Mast/stem cell growth factor receptor KIT, platelet derived growth factor receptor beta (PDGFRβ), anaplastic leukemia kinase (ALK), proto-oncogene tyrosine-protein kinase SRC, tyrosine protein kinase LYN and Bruton’s tyrosine kinase (BTK). Kinases were diluted to approximately 120 nM in kinase dilution buffer (20 mM 4-morpholinepropanesulfonic acid (MOPS) pH 7.5, 1 mM EDTA, 0.01% Brij-35, 5% Glycerol, 0.1% beta-mercaptoethanol and 1 mg/mL bovine serum albumin (BSA). Recombinant kinases (20 nM) were pre-incubated for 15 minutes in reaction buffer containing 25 mM 4-(2-hydroxyethyl)-1-piperazineethanesulfonic acid (HEPES) pH 7.5, 10 mM MgCl_2_, 100 μM ATP, 3 mM DTT, 3 μM NaVO_3_ at 37 °C and 5% DMSO. The assay reaction was started by adding the peptide substrate to a 37.5 μM reaction concentration in a 30 μL volume. Sample aliquots (10 μL) were quenched by combining 1:1 with 30 mM EDTA at 2 and 60-minute timepoints (or 30-minute timepoint for TKI dose response assays).

### Chemifluorescence Detection of Phosphorylation

Quenched aliquots were incubated in a 96-well streptavidin coated plate (125 pmol binding capacity, ThermoScientific) in Tris buffered saline (25 mM Tris-HCL and 150 mM NaCl) with 0.05% Tween 20 (TBS-T) containing 5% w/v skim milk for 1 h.^36^ Subsequently, each well was washed with TBS-T and incubated with antiphosphotyrosine mouse monoclonal antibody 4G10 (MilliporeSigma, 1:10,000 dilution in TBS-T). Following incubation, wells were washed with TBS-T and incubated with horseradish peroxidase-conjugated rabbit anti-mouse immunoglobulin secondary antibody (Abcam, 1:15,000 dilution in TBS-T) for 1h. The wells were washed (TBS-T and 50 mM sodium phosphate buffer) and then treated with Amplex Red (Invitrogen, Carlsbad, CA) reaction buffer (0.5 mM AR, 20 mM H_2_O_2_ and sodium phosphate buffer) for 30 minutes. Fluorescence measurements were taken on a Neo2 microplate reader (Biotek, Winooski, VT) with an excitation wavelength of 530 nm and emission wavelength of 590 nm. The IC_50_ values were calculated by fitting the data to the equation below, were the inhibition_max_ represents the lower plateau of the curve while the inhibition_min_ pertains to the upper plateau, X represents inhibitor concentration and the steepness of the curve was set to a standard hill slope of negative one.

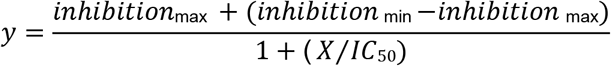

### Experimental design and statistical rationale

The trypsin digestion “library” preparation was performed for two independent replicates, each of which was subjected to an FLT3 kinase variant enzyme in triplicate (Figure S1). Upon file conversion of raw mass spectrometer files into a MGF format, the resulting six kinase treated files per FLT3 variant were combined into one ProteinPilot 5.0 protein identification search. These proteomic database search results were processed as described above (KinaMine and R data formatting scripts) to generate the KINATEST-ID input lists (Figure 1 step 5 and Figure 3).

A Dixon’s Q test with an α value of 0.05 and a ROUT test with a q value set at 5 percent were performed to identify experimental outliers in the inhibitor dose response assay. Fluorescence measurements per well were normalized to the vehicle-only (DMSO) control signal, plotted using Prism software (GraphPad, La Jolla, CA) and fitted to a non-linear equation described above. An exact sum-of-squares F test with a p value set at 0.05 was performed to identify differences in the reported IC_50_ curves for each TKI against the FLT3 kinase variants.

## Results

### *In vitro* kinase reaction to identify substrates for input/analysis with the KINATEST-ID pipeline

Similar to the KALIP method^26^ and others,^37^ we used trypsin-digested cell lysate as a non-randomized peptide “library” to determine FLT3 kinase substrate preferences. Briefly, AML KG-1 cells were grown to log phase, lysed with a urea lysis buffer and digested with trypsin as described above. Following trypsin digest, peptides were treated with alkaline phosphatase to remove endogenous phosphorylation from tyrosines. The phosphatase-treated digest was then divided into aliquots that were processed in parallel: one treated with kinase reaction mixture but no kinase (“FLT3-”), alongside the kinase reaction for each version of the kinase (recombinant FLT3-WT, D835Y or ITD kinase, respectively, “FLT3+”) (Figure 1). The kinase reactions were performed for 16 hours (WT, D835Y and ITD) as described in the original KALIP protocol.^26^ Following kinase treatment, phosphopeptides were enriched using the soluble polyMAC dendrimer^26,27,38^ and analyzed on an Orbitrap Fusion mass spectrometer. Sequence and phosphorylation site identification was performed using the ProteinPilot 5.0 software on the GalaxyP platform (http://galaxyp.msi.umn).

Overall, we identified more than 10-fold more substrates for FLT3 and the two mutant variants than we had curated for most of the kinases we had evaluated in our previous publication using KINATEST-ID, in which substrate numbers ranged from ~15 to ~170.^31^ Using a relatively strict identification quality cut-off of 1% false discovery rate (FDR), we observed 1559 phosphorylated peptides from the kinase reaction with the WT FLT3 treatment, 2010 from the FLT3-D835Y mutant, and 344 from the FLT3-ITD mutant. Of these, 244 overlapped in common from each reaction (Figure 1). The FLT3-D835Y mutation stabilizes the activation loop within the kinase domain, leading to a stable, constitutively active kinase,^5,10^ which likely explains the higher number of phosphopeptides observed in that reaction. The FLT3-ITD mutant is an in-frame gene sequence duplication that encodes for the amino acid segment connecting the juxtamembrane and tyrosine kinase domains. While the WT and D835Y versions of the kinase used in these KALIP experiments contained only a His tag (as described by the vendor), the ITD mutant was also tagged with a GST tag. Due to its larger molecular weight relative to the WT and D835Y variants, less of this kinase was used in the KALIP reactions, which likely accounts for the lower number of substrates observed for that variant. Nevertheless, compared to the manual curation of the literature-reported substrates and phosphoproteomic databases that was implemented previously,^31^ even for the ITD mutant our approach identified many more phosphopeptide sequences than before, which could be used as input for the KINATEST-ID pipeline.

We used either the full list of substrate sequences from each kinase reaction (“WT-16H”, “D835Y-16H” or the “ITD-16H” datasets), or the 244 common substrate sequences that had been identified from all three of the FLT3 variants’ reactions (the “SHARED-16H” dataset), as separate inputs for initial positional scoring matrix (PSM) analyses in KINATEST-ID. The shared substrate sequences were extracted from the phosphoproteomics data outputs using data processing and analysis tools, the KinaMine and Commonality and Difference Finder and the Kinatest part 1.r script (Figure 2), described in the supplementary methods section, to generate PSMs of amino acid preference motifs from each dataset (Figure 3). We observed subtle, but likely functionally insignificant, differences in amino acid over‐ and under-representation at the positions −4 to +4 relative to the phosphorylated tyrosine for each of the different FLT3 variants. Generally for all variants, acidic amino acids were overrepresented and basic amino acids were underrepresented N-terminal to the phosphotyrosine, while hydrophobic amino acids and glutamine/asparagine were slightly overrepresented C-terminal to the phosphotyrosine. Given the lack of substantial differences between the PSMs for the WT and mutant variants, we focused on the substrates that were observed in common for all three kinase variants to move forward with design of novel peptides that could be used as substrates in FLT3 activity assays.

**Figure 2.**
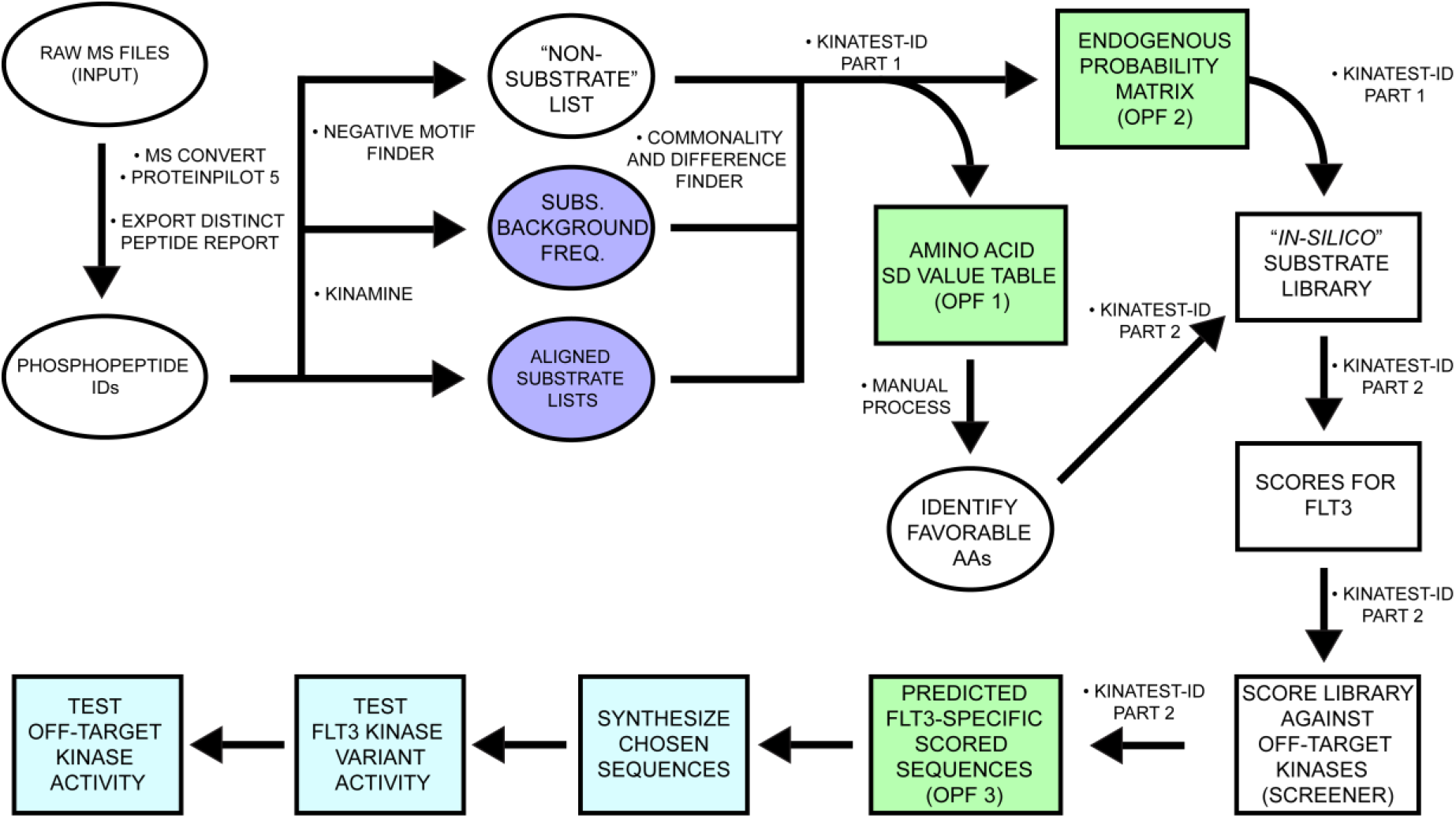
Conceptual overview of our KALIP data processing and formatting for KINATEST-ID incorporation to develop FLT3 artificial substrates. After conversion and peptide/protein ID in the Galaxy-P proteomic pipeline, the KinaMine data formatter tool extracted the confidently-identified (1% FDR) sequences that contained a phosphorylated tyrosine residue (pY), centrally aligned the sequences to the tyrosine of interest (“Aligned substrate lists”) and extracted the UniProt accession number with the accompanying proteins’ amino acid composition file (“Subs. Background Freq.”). The *Negative motif finder* script uses the Uniprot accession numbers to generate a list of tyrosine-containing potential tryptic peptides that were not observed in the phosphopeptide dataset. The scripts KinatestID part1.r and ‐part2.r processed those input files to identify substrate preferences and generate a ranked list of candidate sequences as potential FLT3 substrates. Kinatestid-part2.r then scores the input substrate and non-substrate lists and outputs as two additional files. After performing this workflow on phosphopeptide data from FLT3 kinase reactions, chosen candidate sequences were synthesized and validated *in vitro* against the FLT3 kinase variants. The candidate sequences were then assayed *in vitro* against a panel of kinases to determine off-target kinase activity. *In silico*/predictive steps are illustrated in white/green. Empirical steps of synthesizing peptides and characterizing FLT3 activity and specificity are depicted in light blue.

**Figure 3.**
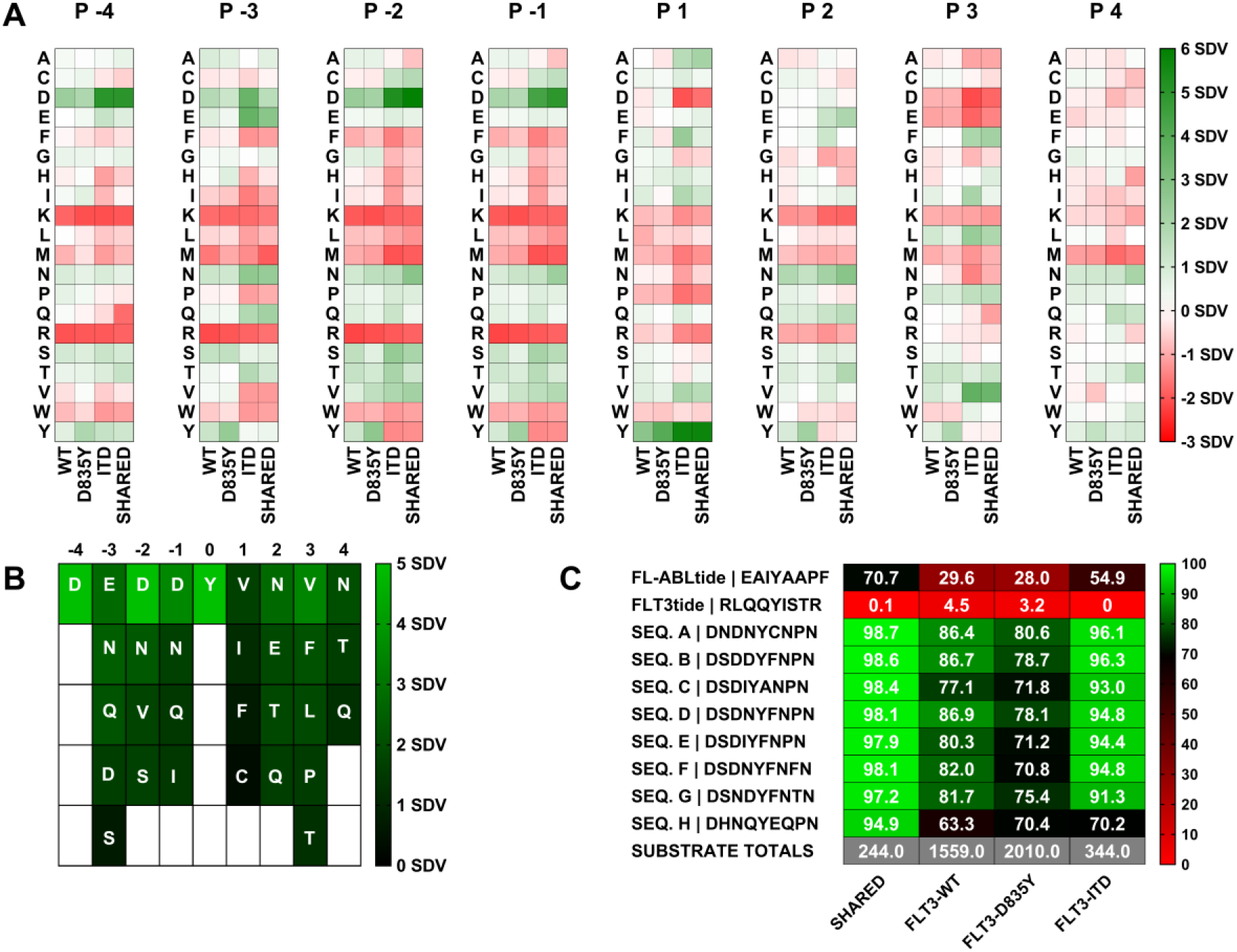
Positional preferences, motif, and substrate candidate mini-library for FLT3-WT, FLT3-D835Y and FLT3-ITD. (A) Observed representation of each amino acid at each position (−4 to +4 relative to phosphotyrosine) in the individual phosphoproteomics datasets for the three FLT3 variants (WT, D835Y, and ITD) and for the sequences shared in all three datasets (Shared). Green = over-represented, white = neutral, red = under-represented. (B) Table summarizing the positional preferences (>1 standard deviation from the mean) used in the “Generator” tool portion of Kinatest part 2.r for permutations to rationally design candidate substrate sequences. Representation of the given amino acids at respective positions is shown via color scale for reference. Sequences of selected, synthesized candidate substrates and controls and scores (illustrated with color scale) against the respective scoring models for each dataset (shared and individual variants).

### KINATEST-ID-based design of novel <u>F</u>LT3 <u>A</u>rtificial <u>S</u>ubstrate pep<u>tides</u> (FAStides)

We then used the “SHARED-16H” dataset and employed the next steps of the KINATEST-ID approach to design a set of candidate sequences for synthesis and biochemical testing. This process used the KINATEST-ID “Generator” tool (via the Kinatest part 2.r script) to create a list of sequences comprising all the permutations of the amino acids overrepresented at each position by at least two standard deviations from the mean (Figure 3A-B). One caveat was that while tyrosine was observed as overrepresented at −1 and +1 to the phosphotyrosine, we chose to exclude it from the preference motif for this iteration of substrate design, due to the potential ambiguity in assay signal that could ultimately be introduced by having more than one phosphorylatable residue in the designed substrates. Additionally, F was included as an option at position +1 (despite having relatively low representation) as a hydrophobic alternative to the more highly represented I and V in an attempt to provide better specificity, due to the high frequency of those two amino acids at that position in the motifs for other tyrosine kinases. Permutation of the motif was then followed by scoring of the resulting 19,201 sequences against the PSMs for WT FLT3, the two mutant variants, and a panel of other kinases,^31^ using the KINATEST-ID “Screener” tool (Figure 2). The sequences and their scores against the PSMs are summarized in Figure 3C while their “off-target” kinase PSM scores are summarized in Figure 5G. We chose one set of sequences that scored well for FLT3 but scored poorly for other kinases (sequences A, D, G and H). Since the highest scoring sequences for FLT3 also scored well for several other off-target kinases in the Screener panel, we selected another set of those sequences (sequences B, C, E and F) to have a higher likelihood of obtaining an efficient (though potentially not FLT3-specific) substrate. Additionally, we synthesized two control sequences (Tables 1 and 2) that were previously reported to be phosphorylated by FLT3 (ABLtide and FLT3tide).^39,40^

**Table 1.** IC_50_ values measured by monitoring the phosphorylation of FAStide-E or ‐F in ELISA-based assays.

| Table 1: TKI IC<sub>50</sub> VALUES (pM) +/- S E |  |  |  |  |  |  |
| --- | --- | --- | --- | --- | --- | --- |
| TKI | FLT3-WT |  | FLT3-ITD |  | FLT3-D835Y |  |
|  | Seq. E | Seq. F | Seq. E | Seq. F | Seq. E | Seq. F |
| SORAFENIB | 8.7±3.5 | 13±11 | 780±300 | 550±81 | 1,800±300 | 3,200±940 |
| QUIZARTINIB | 8.6±1.3 | 32±20 | 190 ±110 | 266 ±49 | NA | 4,700±870 |
| CRENOLANIB | 13±3.8 | 1.3±0.92 | 45 ±14 | 52 ±8.9 | 15±1.4 | 32±7.9 |

### *In vitro* validation of FAStide sequences as FLT3 kinase variant substrates

We tested the candidate sequences via *in vitro* kinase assays using anti-phosphotyrosine antibody 4G10 in a chemifluorescent Enzyme-Linked Immunosorbent assay (ELISA) as the readout to detect peptide phosphorylation, with sample aliquots quenched at 2 and 60 minutes. Figure 4A shows that the sequences that were most efficiently phosphorylated by FLT3-WT, D835Y and ITD contained the DXDXYXNXN motif. Figure 4A shows that S was well-tolerated at position −3 (while N and H were less tolerated), both N and D were tolerated at position −1, and F was the preferred amino acid at position +1 while A was not well tolerated. Sequences that contained F, P or T residues at position +3 were phosphorylated by all FLT3 kinases. The control peptides (FL-ABLtide and FLT3tide) were also scored against the PSMs for all of the datasets (Figure 3C) and assayed in parallel with the FAStide sequences against the FLT3 kinase variants. The previously reported substrate FLT3tide^39^ scored poorly against our models and was a poor FLT3 substrate in our assays (Figure 4A). ABLtide, a previously reported FLT3 substrate,^40^ scored moderately against the SHARED-16H dataset model, and performed moderately as a substrate.

**Figure 4.**
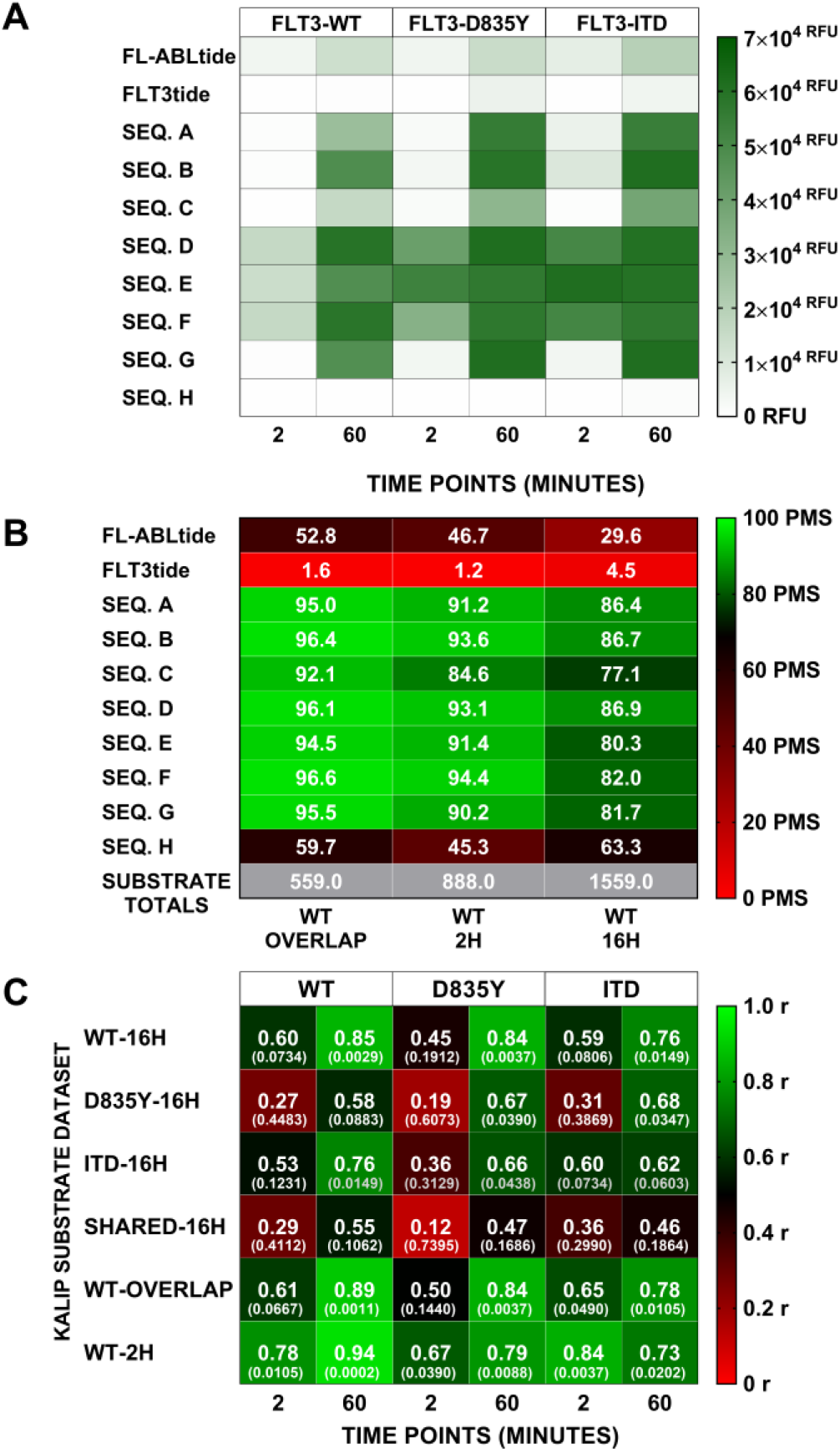
FLT3 kinase variant activity assays and scoring model correlation. (A) FLT3 (WT, D835Y and ITD) in vitro kinase assay results for the candidate sequences (A, B, C, D, E, F, G and H). Each FLT3 variant (columns WT, D835Y, ITD) was reacted with each peptide (rows), with aliquots taken at 2 and 60 min (sub-columns). Phosphorylation levels indicated by RFU from ELISA detection, illustrated via color scale (green = high phosphorylation, white = low/no phosphorylation). (B) Effect of *in vitro* KALIP kinase assay incubation time on KINATEST-ID scores for candidate sequences. Positional scoring model scores for each sequence (rows) against the models (columns) derived from all phosphopeptides observed in the WT 2H or 16H KALIP kinase reaction, respectively, or just those observed in both (WT OVERLAP). Color scale from red (low PMS score) to green (high PMS score). (C) Spearman’s non-parametric correlation test to measure correlation between the KINATEST-ID’s scoring system and the *in vitro* activity assay results. r values shown in bold, above p-values denoted in parenthesis. Color scale indicates low (red) to high (green) Spearman r values.

### Evaluating the relationship between substrate input datasets and resulting PSM model scores vs. biochemical assays

FAStide sequences A, B, D, E, F and G generally had higher PSM scores than C and H, and were all phosphorylated more efficiently than C and H in the assays. The FLT3tide reference peptide scored poorly in all the matrices and was phosphorylated very poorly in the assays, and FL-ABLtide scored moderately and was phosphorylated moderately relative to the best of the FAStide sequences in the assays. For the longer KALIP kinase reactions (16 hours), the strongest correlations were found between each variant’s biochemical assay results at 60 min and the WT-16H dataset-derived PSM scores (Figure 3C). The FAStide sequences scored the lowest with the D835Y-16H PSM model, which contained the largest substrate list (2010 substrates), and their scores had lower Spearman correlations with assay results from the WT, D835Y, and ITD variants, respectively. This is most likely attributable to a combination of the larger size of the input dataset for that PSM and that the FAStides were designed based on the selected subset of that data shared with the other two variants, rather than the entire dataset used to derive this PSM model. The FAStide sequences received higher scores using PSM models with the smallest substrate lists (SHARED-16H and ITD-16H), which was also primarily an artifact of all or nearly all of the sequences in those smaller datasets being used to design the FAStides in the first place. However those scores were less correlated with the biochemical assay results, which might also arise from smaller dataset artifacts or some other, as yet unidentified factor affecting the accuracy of the predictive models from those datasets.

### Evaluating the length of time of *in vitro* kinase treatment on substrate motif prediction

We also examined the effect of reaction time in the kinase treatment step on the ability of the KALIP-KINATEST-ID process to identify efficient substrates that can be used in enzyme assays. We performed a two-hour kinase reaction using FLT3-WT kinase and processed the data (referred to hereafter as WT-2H) as described above to determine if the KALIP kinase treatment time affected 1) the characteristics of the preference motif arising from a given dataset, and 2) its utility for subsequent substrate design. We identified 888 phosphopeptides from the 2H KALIP FLT3-WT kinase treatment (relative to 1559 for the 16-H treatment as described above). We compared the “WT-2H” to the “WT-16H” substrate list and found 559 sequences shared by both. These sequences, referred to as the “WT-OVERLAP” substrate and background frequency lists, represented sequences that were likely to have been phosphorylated rapidly and robustly (Figure 4B). The corresponding SDV values were compared to those for each substrate list from the WT-2H and WT-16H experiments, as shown in Figure S3.

Overall, the preferences at each position as represented by the SDV tables were similar (Figure S3), however subtle differences in the WT-2H dataset from the two hour incubation resulted in a scoring model that appeared to more accurately reflect the substrate phosphorylation efficiency for the WT kinase as observed in the biochemical experiments relative to the models derived from the WT-16H dataset. PSM scores derived from the WT-2H, WT-16H and WT OVERLAP datasets were compared with the assay results for the eight FAStides and two control peptides (Figure 4A). Spearman correlations are shown in Figure 4C. Similar to the WT-16H, the assay signal at 60 min and the PSM scores generated via the WT-2H and WT-OVERLAP datasets were very highly correlated, appearing slightly stronger for WT-2H than for WT-OVERLAP and WT 16H. This may indicate that shorter KALIP kinase treatment time is better for determining more efficiently (i.e. rapidly) phosphorylated substrates, and that longer treatment may “dilute” the substrate preference motif with less efficient sequences.

### *In vitro* characterization of FAStide FLT3 specificity

While *in vitro* recombinant kinase assays using the novel peptides identified here would not require exquisite specificity since they would typically employ purified kinase, we wanted to perform a limited assessment of the “off-target” phosphorylation of these peptides using a small panel of other kinases: KIT, PDGFRβ, ALK, SRC, LYN and BTK. KIT and PDGFRβ are kinases for which the substrate preference motif has not yet been identified, however, they are within the same kinase family as FLT3 and based on observations for other kinases,^31^ were likely to phosphorylate the same FAStides as FLT3. Based on the previously published PSM scoring models^31^ (derived from manual substrate list curation and not the KALIP process), SRC was predicted to phosphorylate sequences B, C, E and F while LYN is predicted to phosphorylate sequences A and C. BTK was not expected to phosphorylate any of the FAStide sequences. Additionally, our panel included the receptor tyrosine kinase ALK, which we previously observed to phosphorylate similar sequences to those phosphorylated by SRC.^41^ To ensure each of those recombinant kinases was active, we performed an *in vitro* kinase assay with several reference peptides that have been previously characterized in our laboratory for the kinases in the panel. The “universal” tyrosine kinase substrate that we previously reported U5 (DEAIYATVA)^34^ was the reference peptide chosen for KIT, PDGFRβ and BTK, and an ALK substrate we previously reported (ALAStide)^41,42^ was used for ALK (Figure 5A-F). SFAStide-A (DEDIYEELD)^31^ was used as the reference peptide for SRC and LYN kinases. Briefly, the kinases were preincubated with kinase reaction mixture and the *in vitro* reaction was initiated by the addition of substrate peptide. The samples were quenched at 2 and 60-minute time points as described above. Phosphorylation was measured using the previously described ELISA-based assay^31,36^ and results are shown in Figure 5.

**Figure 5.**
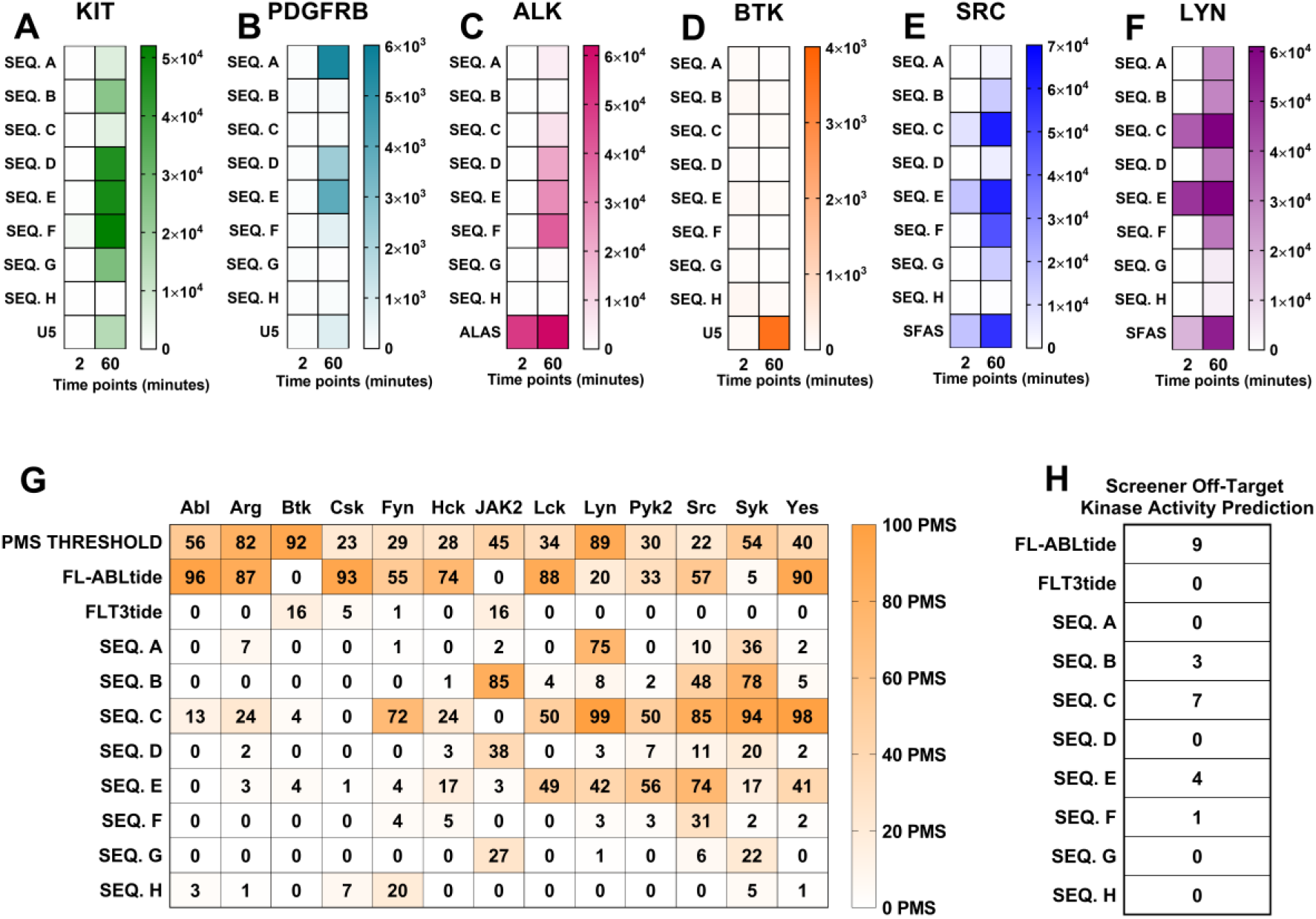
The focused peptide library was assayed against a panel of kinases in a limited test of FLT3 specificity. (A-F) Phosphorylation level for each peptide (rows) at two time points (2 and 60 min, columns) is shown separately for each kinase, with color scales normalized to minimum/maximum RFU values from ELISA detection for that kinase. FAStide candidate data shown in rows 1-8, reference peptide data show in row 9 as a positive control for kinase activity. (G) PMS scores for each FAStide candidate and two reference peptides (rows) for 13 kinases (columns) previously built into the KINATEST-ID Screener tool from literature curated substrate input lists. Scores are indicated by color scale from low (white) to high (orange). (H) “Off-target” activity prediction from Screener tool, indicating the number of kinases predicted to tolerate a given sequence as a substrate.

Overall, the sequences with the least off-target phosphorylation in this panel were FAStide-A, which was phosphorylated moderately by c-KIT and PDGFR**β** (both FLT3 family members) and LYN after 60 minutes but not the others in the panel, and FAStide-G, which was phosphorylated to a relatively low degree after 60 minutes only by c-KIT and SRC (Figure 5A and 5E). FAStide-B, ‐D, ‐E and ‐F were phosphorylated by more of the off-target kinases by 60 min and FAStide-C was a robust substrate for both SRC and LYN. PDGFRβ and BTK, on the other hand, did not phosphorylate any of the artificial sequences over a 60-minute incubation. The off-target *in vitro* kinase assay results were mostly but not entirely consistent with the Screener predictions, which is not surprising given that Screener is limited by the PSM models built into its cross-referencing algorithm—the main caveat is that the PSM models in Screener all come from the previously developed KINATEST-ID package^31^ that did not have KALIP phosphoproteomics data as input.A future goal is to update current kinase PSMs with newly generated KALIP data, as well as adding data from more kinases to improve the cross-referencing depth.

### Detection of FLT3 kinase variant inhibition through FAStide *in vitro* phosphorylation

To demonstrate how the FLT3 artificial substrates can be used to monitor TKI efficacy, we performed dose-response (DR) assays for FLT3-WT, FLT3-D835Y and FLT3-ITD with sorafenib, quizartinib and crenolanib, three TKIs that have been characterized against the three FLT3 variants.^20,22,25,43^ FAStide-E and FAStide–F were chosen for the DR assays due to their efficient phosphorylation by all three FLT3 kinase variants, and employed in parallel experiments. Each FLT3 kinase variant was pre-incubated in the kinase reaction mixture (containing ATP) with the respective TKI (0.00001 to 100 nM) for 15 minutes at 37°C without substrate, and the kinase reaction was initiated via the addition of the substrate (37.5 μM). Reactions were quenched after 30 min and wells analyzed using ELISA as described above. Fluorescence values (relative fluorescence units, RFU) were collected and normalized to values for wells containing vehicle control (DMSO). In general, both substrates exhibited dose-response curves and IC_50_ values that were consistent with what was expected for the given inhibitor against each FLT3 variant, with one notable exception (further described below) (Figure 6, Table 1). All three inhibitors potently inhibited WT FLT3 (with IC_50_ values in the ~1-30 pM range). Potency towards the ITD mutant was lower for all three inhibitors (IC_50_ values between ~40-800 pM), with crenolanib more potent than the other two. The D835Y mutant’s dose response curves were also as expected for sorafenib and crenolanib, with sorafenib being significantly less potent than it was against the WT (~200-250-fold higher IC_50_), while crenolanib maintained pM-range IC_50_. For quizartinib, on the other hand, dose response curves were different for the assays performed using FAStide-E compared to FAStide-F. Quizartinib is a type II inhibitor, known to bind to the inactive “DFG-out” conformation of the kinase. Bulky, hydrophobic mutations at position 835 in FLT3 are thought to confer resistance to quizartinib, due to the effects of the side chains on the structure and dynamics of the DFG loop in the kinase domain, with the extra steric bulk disrupting the stability of the inactive conformation.^44,45^ The FAStide-E quizartinib dose response results for the D835Y mutant were consistent with this model, with essentially no significant inhibition even at concentrations as high as 100 nM (>10,000-fold higher than the IC_50_ for quizartinib against the WT FLT3). However using FAStide-F as the substrate, inhibition was observed in the same IC_50_ range as for sorafenib. This suggests that substrate interactions may affect inhibitor binding stability, perhaps by playing a role in DFG-in/-out dynamics.

**Figure 6.**
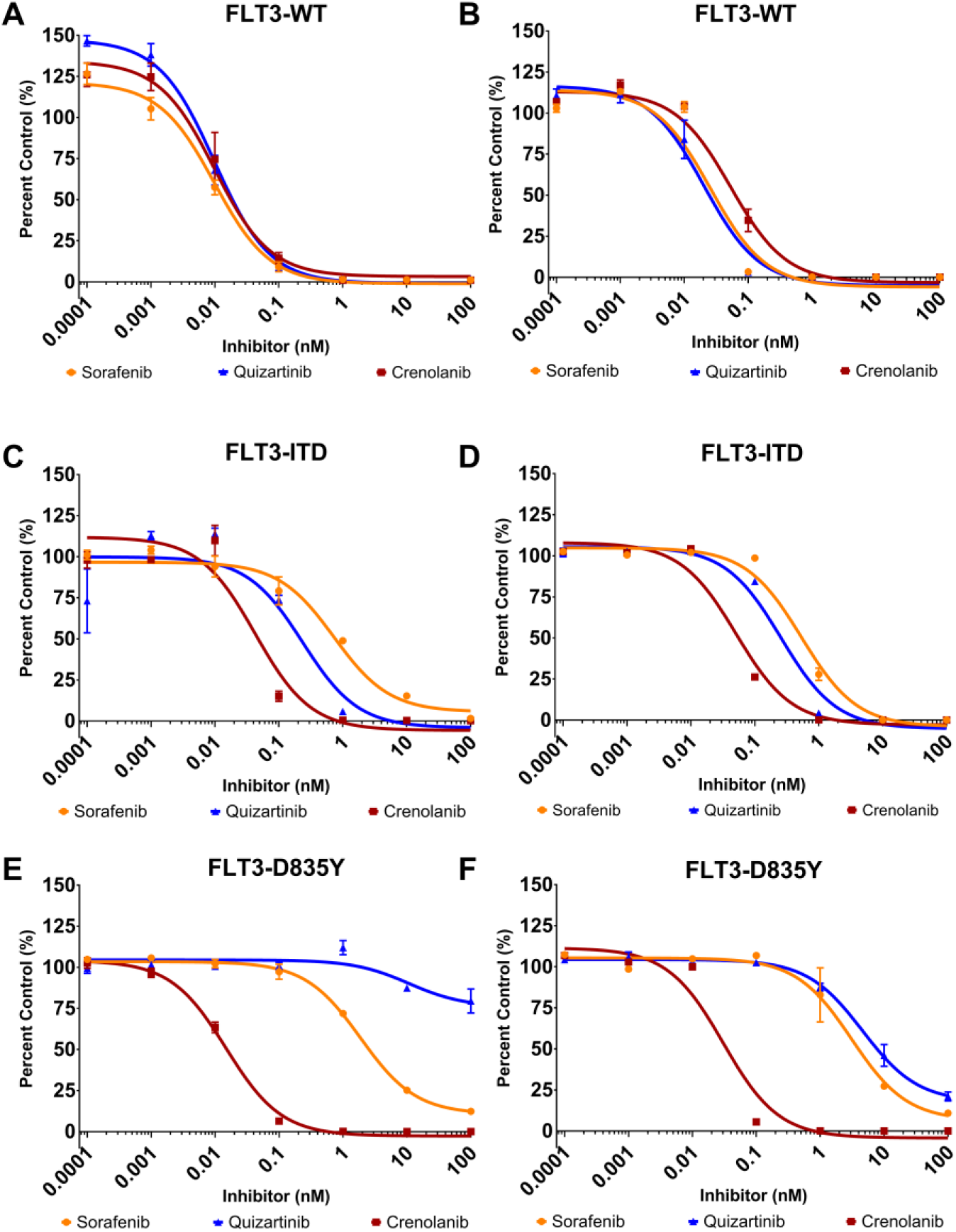
Monitoring FLT3 kinase activity and inhibition by clinically relevant tyrosine kinase inhibitors (TKI). Dose-response assays were performed using two substrates, FAStide-E (A, C, E) and FAStide-F (B, D, F) in the presence of increasing TKI concentrations (sorafenib, quizartinib and crenolanib). Data show RFU values for each inhibitor normalized to vehicle control (DMSO; representative of 6 independent reactions) and plotted as percent control. IC_50_ values were calculated from fitting the data to a fixed slope (three parameter) curve in GraphPad Prism. Data points/error bars represent mean ±SEM.

## Discussion

Drug resistance in AML has been a major factor in the poor 5-year survival and clinical remission rates. Treatments targeting patients with FLT3-positive AML have seen promising results, but inhibitor resistance has been detrimental to clinical efficacy. FLT3 remains a viable drug target in AML,^40,46^ however, none of the current TKIs used in therapy are FLT3 specific and even once those are developed, rapid emergence of mutations that abrogate drug binding will be a continuing challenge.^11,12,47,48^ This highlights the importance of having efficient assays that can be used as tools to identify specific and selective TKIs that target FLT3 and mutant variants. In this work, we coupled *in vitro* kinase reactions on cell lysate digests with the KINATEST-ID pipeline to design, synthesize and validate a panel of sequences to detect the activity of a kinase that has few known substrates. Our process created a novel panel of peptides that can be used in kinase assays and provide higher phosphorylation efficiency than previously reported substrates. These findings demonstrate how the streamlined combination of KALIP and the KINATEST-ID pipeline can be used to identify novel artificial kinase substrates.

The original KINATEST-ID pipeline^31^ relied on literature-validated sequences, including some from proteomic databases and positional scanning peptide library (PSPL) assays, as input for the matrices. Each sequence was manually curated by further literature examination (looking for corroboration of upstream kinase evidence via e.g. testing of non-phosphorylatable A/F mutations). This severely limited the number of substrates that could be included in the “true positive” input list, which likely resulted in less accurate predictions of optimal substrate sequences. While this was sufficient to develop effective substrates for several kinases that showed reasonable degrees of selectivity for their targets,^31^ it was not optimal—and further, if a given kinase did not have sufficient known substrates or PSPL assay data available then it was not possible to make any prediction at all. Using cell lysate digest as a peptide library for high-throughput identification of FLT3 kinase variant substrates enabled a large increase in the numbers of *bona fide* substrate sequences that could be used to build positional scoring models.

The improved positional scoring matrix models developed from these large, empirically detected substrate sequence datasets enabled a prediction for FLT3’s preferred amino acid motif, which was then used to design several potential novel substrate peptides. Scores from the positional scoring matrix models correlated well with the relative biochemical behavior of the novel substrates, especially when the input dataset comprised sequences that were observed as phosphorylated after a short kinase incubation time (2 h, which is closer to the reaction time scale used for biochemical assays in practice). This suggests that even though endogenously-derived tryptic peptide libraries are somewhat biased relative to randomized/unbiased synthetic libraries^29^ (given the sequence constraints imposed by their genomic origin), they are still able to provide sufficient sequence diversity to enable discovery of hundreds to thousands of substrates and accurately reveal substrate preferences for a given kinase. It also suggests that although performing the reaction at the protein level^30^ may be better for developing prediction models for identifying endogenous protein substrates, performing substrate preference analyses at the peptide level *in vitro* is sufficient for designing peptide probes.

Intriguingly, we observed substrate-dependent inhibition for the well-characterized TKI quizartinib against the FLT3 D835Y mutant. This suggests that the particular substrate used in a screening assay might bias the interpretation of whether an inhibitor is or is not potent against a given enzyme. This highlights the importance of the substrate in inhibitor assays, and suggests that expanding the range of efficient substrates available as drug discovery tools would be beneficial. Work is ongoing to determine whether this is specific to the FLT3 D835Y mutant or is a more general issue for kinases. Other next steps will be to expand the application of this approach to the kinases previously built into the KINATEST-ID “Screener” panel,^31^ in order to improve the accuracy of the positional scoring matrix models for the “off-target” kinases and achieve better predictions of selectivity during the substrate design process. While the original, previously published Screener panel was accurate enough to offer the practical ability to pre-filter a large list of potential sequences down to a more manageable number, clearly the selectivity prediction was limited by the same factors (comprehensiveness of the input dataset) as the preference prediction. Ongoing efforts to apply the KALIP adaptation approach reported here to more kinases should facilitate improvement of this aspect, as well.

In summary, in this work we demonstrate the utility of generating a large dataset of *bona fide* substrate information, using a relatively cheap and easily produced peptide pseudo-”library” derived from cellular proteins via proteolytic digest, for defining substrate preference motifs and scoring models that enable design of efficient peptide substrates for kinase enzymes. This strategy enabled discovery of multiple substrates, some of which may influence inhibitor interactions with the enzyme and affect conclusions about inhibitor efficacy. We also anticipate that this process can be applied to orphan kinases for which little to nothing is known, to first identify substrate sequences through the *in vitro* proteolytic peptide library kinase reaction followed by prediction of the preference motifs from those data. Those motifs could be used to design, synthesize and validate artificial substrates, which can assist in chemical biology and drug discovery efforts to identify novel and potent inhibitors to study their biology and/or become therapeutic leads. Furthermore, this workflow could potentially be generalized to any enzyme driven disease for which substrate preference data can be determined from proteolytically or synthetically prepared peptide libraries^49^ and used to design novel substrates for use in assays. This will greatly enhance the generalization of the novel substrate probe design process we initially implemented in our first report of KINATEST-ID,^31^ broadening the scope for drug discovery assay development.

## Acknowledgments

This work was supported by the National Cancer institute, Center to Reduce Cancer Health Disparities Diversity Training Branch: R33CA183671, R01CA182543 and R01CA182543-03S1 We thank Dr. W. Andy Tao (Department of Medicinal Chemistry and Molecular Pharmacology, Purdue University) for his helpful discussions and insights in adapting the kinase assay linked with phosphoproteomics technique. We thank the University of Minnesota’s Center for Mass Spectrometry and Proteomics for assistance in data acquisition and analysis. We thank Dr. Anton B. Iliuk (Tymora Analytica, West Lafayette, IN) for his support and insight in implementing the PolyMAC phosphopeptide enrichment kit.

**Abstract Figure 1.**
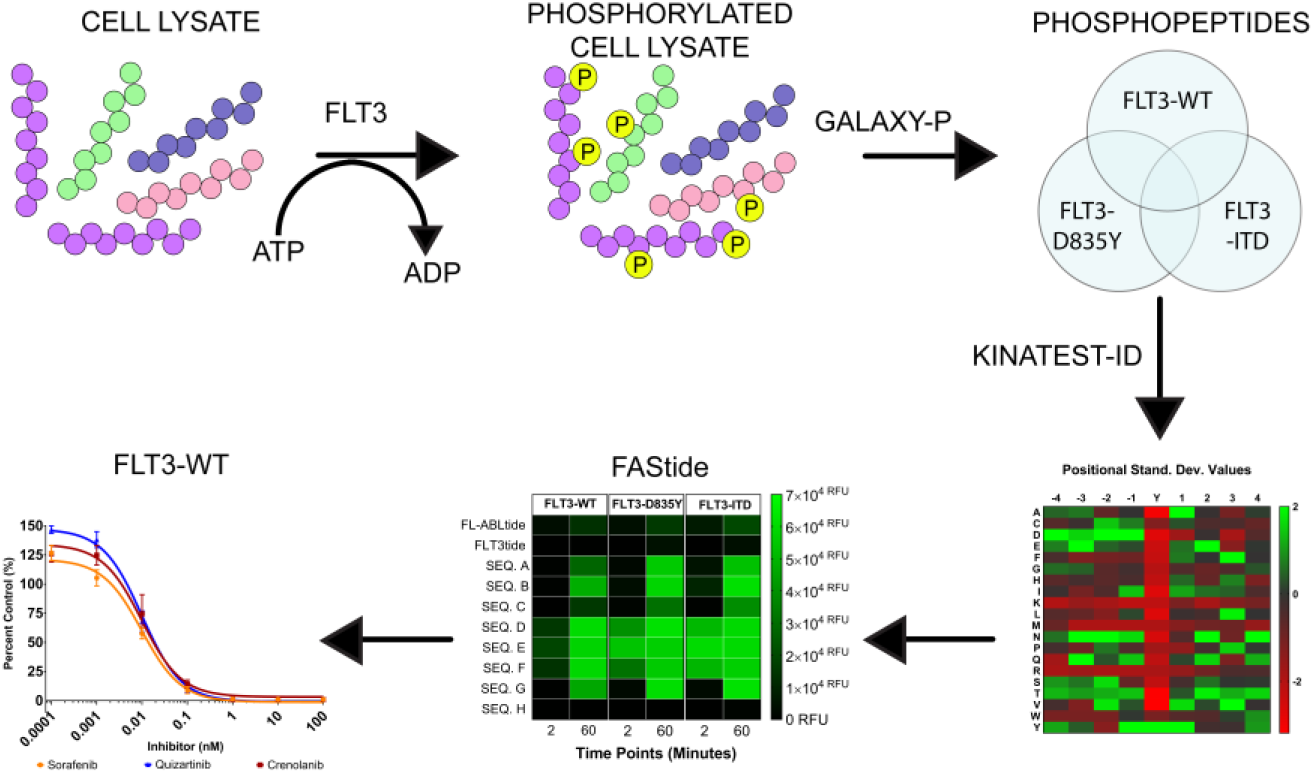

## Data availability

The data is accessible at PRIDE.

## List of abbreviations

AML: acute myeloid leukemia
RTK: receptor tyrosine kinase
ITD: internal tandem duplication
TKD: tyrosine kinase domain
WT: wild type
TKI: tyrosine kinase inhibitor
KALIP: Kinase Assay Linked Phosphoproteomics
FBS: fetal bovine serum
PBS: phosphate buffered saline
ACN: acetonitrile
DTT: dithiothreitol
TFA: trifluoroacetic acid
HLB: hydrophilic-lipophilic balanced
Tris-HCL: tris(hydroxymethyl)aminomethane hydrochloride
EDTA: ethylenediaminetetraacetic acid
FA: formic acid
FDR: false discovery rate
MCC: Matthews Correlation Coefficient
AA: amino acid
DMF: dimethylformamide
DMSO: dimethyl sulfoxide
PDGFRβ: platelet derived growth factor beta
ALK: anaplastic leukemia kinase
BTK: burton’s tyrosine kinase
MOPS: 4-morpholinepropanesulfonic acid
BSA: bovine serum albumin
HEPES: 4-(2-hydroxyethyl)-1-piperzoneethanesulfonic acid
TBS: tris buffered saline
AR: amplex red
2H: two hours
16H: 16 hours
SDV: standard deviation values
PPM: positional probability matrix
SSM: site selectivity matrix
DR: dose response

